# CRISPR-interference based modulation of mobile genetic elements in bacteria

**DOI:** 10.1101/428029

**Authors:** Ákos Nyerges, Balázs Bálint, Judit Cseklye, István Nagy, Csaba Pál, Tamás Fehér

## Abstract

Spontaneous mutagenesis of synthetic genetic constructs by mobile genetic elements frequently results in the rapid loss of advantageous functions. Previous efforts to minimize such mutations required the exceedingly time-consuming manipulation of bacterial chromosomes and the complete removal of insertional sequences (ISes). To this aim, we developed a single plasmid-based system (pCRIS) that applies CRISPR-interference to inhibit the transposition of bacterial ISes. pCRIS expresses multiple guide RNAs to direct inactivated Cas9 (dCas9) to simultaneously silence IS*1*, IS*3*, IS*5*, and IS*150* at up to 38 chromosomal loci in *Escherichia coli*, *in vivo*. As a result, the transposition rate of all four targeted ISes dropped to negligible levels at both chromosomal and episomal targets, increasing the half-life of exogenous protein expression. Most notably, pCRIS, while requiring only a single plasmid delivery performed within a single day, provided a reduction of IS-mobility comparable to that seen in genome-scale chromosome engineering projects. Global transcriptomics analysis revealed nevertheless only minute alterations in the expression of untargeted genes. Finally, the transposition-silencing effect of pCRIS was easily transferable across multiple *E. coli* strains. The plasticity and robustness of our IS-silencing system make it a promising tool to stabilize bacterial genomes for synthetic biology and industrial biotechnology applications.

## INTRODUCTION

*De novo* constructed DNA elements propagated in living cells frequently impede cellular fitness. In turn, evolutionary processes that are inactivating these DNA elements often release cells from growth retardation and thus lead to the replacement of the engineered population. Therefore these evolutionary processes, while resolving collateral burden, are hindering the central aim of biotechnology and synthetic biology: the predictable engineering of biological systems *via* the introduction of *de novo* constructed DNA elements (1).

Several strategies are already available to deal with this issue (2,3): One possibility is to engineer redundant genetic circuits that can withstand mutations without loss-of-function (4). Another solution is to design DNA sequences that are less prone to mutation-acquisition, mainly by relying on experimental observations and computational designs (5,6). Yet another way to limit the evolution of genes of interest is to link their expression to that of an essential gene (6,7). Finally, the ultimate strategy to mitigate this issue is the reduction of the mutation rate of the engineered host. However, this latter solution until now remained limited to certain mutation types and a few bacterial models. For example, decreasing the rate of point mutations in bacterial cells has been achieved through the overexpression of individual genes (*mutL* (8) or *nudG* (9)), by removing the genes of error-prone DNA polymerases, (*polB, dinB* and *umuDC*) (10), or by mutating the gene encoding RNAse E (11). Also, the removal of the *recA* recombinase (12) or the *dam* methylase (13) have been shown to reduce the frequency of deletions, especially those arising by the homologous recombination of repetitive segments.

Strikingly, a recent study highlighted that insertional mutagenesis, caused primarily by insertion sequences (ISes) constitutes the main cause of instability of synthetic DNA elements in *Escherichia coli (14)*. Bacterial ISes are most probably the simplest autonomous mobile genetic elements found in nature: they consist of a transposase gene (often split into two open reading frames) flanked by terminal inverted repeats (15). ISes have been known to make a measurable contribution to the spontaneous mutation rate of bacterial genomes, accounting for 3.9 to 98 % of genetic changes (16–22). Therefore, genetic stabilization of bacterial hosts must include the complete elimination of their mobility. Based on this concept, the deletion of all transposable elements from the genomes of various *E. coli* strains (23–25) and *Acinetobacter baylyi* (26) have already proven to be a successful strategy to increase the stability of chromosomal or plasmid-based synthetic constructs. One of these endeavors, which removed all mobile DNA (prophages, cryptic prophages, ISes) from the chromosome of *E. coli* MG1655 by subsequently executing 42 precise deletions, took several years to accomplish in our laboratory (23). Later, the availability of programmable CRISPR/Cas (*c*lustered *r*egularly *i*nterspaced *s*hort *p*alindromic *r*epeats and *C*RISPR-*as*sociated proteins) (27,28) endonucleases revolutionized the genome-wide inactivation of mobile DNA both in bacteria (24) and in mammalian cells (29,30). Furthermore, an *in vivo* targeted base-editing technique has been recently developed that could potentially inactivate mobile DNA elements in *E. coli* using a CRISPR/Cas-targeted cytidine deaminase (31).

Recently, the discovery of transcriptional control by CRISPR/Cas interference (CRISPRi) has introduced a novel set of tools for the transcriptional reprogramming of living cells (32–34). Essentially, this system directs a catalytically inactive “dead” Cas nuclease (dCas9) *in vivo* to virtually any target DNA specified by a short complementary RNA molecule (crRNA) and thereby achieves transcriptional-suppression of the target. Due to its robustness and portability, CRISPRi presents a versatile solution for genome-scale control of cellular phenotypes. By building on these advantageous properties, we present a system of plasmid-based CRISPR-interference termed pCRIS to simultaneously inhibit the major sources of insertional mutagenesis in various strains of *E. coli*. We demonstrate the ability of pCRIS to repress multiple bacterial transposases and consequently increase the genetic stability measured at both chromosomal and plasmid-borne loci.

## METHODS AND MATERIALS

### Strains, chemicals, and media

Plasmids were constructed in *Escherichia coli* strain MDS42 (23). Chromosomal mutation rates were measured in *E. coli* strains MG1655 (35) and BL21(DE3) (36). Plasmid stability was monitored in *E. coli* DH5αZ1 (37) and JM107MA2 (38). Bacteria were grown in Luria-Bertani (LB) medium (39) or in mineral salts (MS) minimal medium, supplemented with 0.2 % glucose (17).

Antibiotics (Sigma-Aldrich, St. Louis, MO, USA) were used in the following end-concentrations: chloramphenicol (Cm): 25 μg/ml, ampicillin (Ap): 100 μg/ml, kanamycin (Km): 25 μg/ml, cycloserine (Cyc): 0.04 mM. IPTG was applied in 1 mM end concentration. Standard protocols were followed in all DNA manipulation and cloning processes unless otherwise specified (39).

### CRISPR spacer design

CRISPR spacers were designed to bind the -10 box of targeted promoters. To define the -10 region of the transposase promoters, we analysed the hypothetical promoter-region of each IS element using the Neural Network Promoter Prediction interface (40). The -10 box of IS*5* has been experimentally determined before (41). For IS*150*, we targeted the -10 box of Promoter 1, published earlier (42). The promoter sequences were used as inputs at the Cas Online Designer (http://cas9.wicp.net) to obtain potential spacer sequences. The output CRISPR spacers were manually selected for complete overlap with the predicted -10 box. The Cas Online Designer was used to verify that the chosen spacers targeted all copies of an IS type within the *E. coli* genome, with no additional targets. Each spacer sequence was designed following a previous guideline (34). A dsDNA segment containing a triple CRISPR array targeting IS*1*, IS*5* and IS*3* was synthesized by GenScript (Piscataway, NJ, USA). The two complementary strands of the IS*150*-specific spacer/repeat tandem were synthesized as oligonucleotides. Both the dsDNA and the oligonucleotides were supplemented with appropriate extensions to allow their cloning into the BsaI site of the pCRISPathBrick plasmid or its derivatives (see Table S1).

### Plasmids

Plasmid pdCas9 (Addgene #46569) was a kind gift of Prof. Luciano Marraffini, obtained via Addgene (33). Plasmid pCRISPathBrick was constructed by modifying pdCas9 as described earlier (34). pCRISP_IS was constructed by cloning the triple CRISPR array targeting IS*1*, IS*5*, and IS*3* into the BsaI site of pCRISPathBrick, following the protocol of Cress et al (34). To obtain pCRIS, the two complementary oligonucleotides encoding the IS*150*-specific CRISPR spacer/repeat tandem were annealed and cloned into the BsaI site of pCRISP_IS. Plasmid pBDP_RFP_GFP, harboring a red- and a green fluorescent protein gene, driven by the bi-directional promoter BDP01 was a kind gift of Prof. Herbert M. Sauro (7). Plasmid pBDP_Km_GFP5 was constructed by replacing the *rfp* gene of pBDP_RFP_GFP with the kanamycin resistance (*km*) gene of plasmid pSG76-K (GenBank Acc. No. Y09894.1), as follows: The pBDP_RFP_GFP plasmid, less the *rfp* gene was PCR-amplified using primers pBDPfw and pBDPrev using Dream Taq DNA Polymerase (Thermo Fisher Scientific, Waltham, MA, USA), while the *km* gene of pSG76-K was amplified with primers Km5 and Km3. The two PCR products were gel-purified with the Viogene Gel/PCR DNA Isolation Kit, (Viogene-Biotek Corporation, New Taipei, Taiwan R.O.C.) and fused using the NEBuilder kit (New England Biolabs, Ipswitch, MA, USA), according to the manufacturers’ instruction. The obtained plasmid, pBDP_Km_GFP5 constitutively expresses Ap-resistance and has an IPTG-inducible bidirectional promoter controlling the expression of Km-resistance and GFP. All plasmids were purified using the Zippy Plasmid Mini Prep Kit (Zymo Research Ltd., Orange County, CA, USA)

### RNA preparation and sequencing

Three parallel cultures of *E. coli* strain MG1655 carrying the control pCRISPathBrick plasmid and three cultures of MG1655 carrying the pCRIS plasmid were grown at 37 °C in MS medium supplemented with 0.2 % glucose and Cm. When reaching an OD540 value of 1, each culture was treated with the RNAprotect Bacteria Reagent (Qiagen, Leipzig, Germany), and cellular RNA was prepared using the E.Z.N.A. Bacteria RNA Kit (Omega bio-tek, Norcross, GA, USA) following instructions of the manufacturers. Whole transcriptome sequencing was performed using TrueSeq RNA Library Preparation Kit v2 (Illumina, San Diego, CA, USA) according to the manufacturer’s instructions with slight modifications. Briefly, RNA quality and quantity measurements were performed using RNA ScreenTape and Reagents on TapeStation (all from Agilent, Santa Clara, CA, USA) and Qubit (Thermo Fisher Scientific, Waltham, MA, USA); only high quality (RIN >8.0) total RNA samples were processed. Next, 1 µg of RNA was DNaseI (Thermo Fisher Scientific, Waltham, MA, USA) treated, the ribosomal RNA depleted using RiboZero Magnetic Kit for Gram-negative bacteria (Epicentre, Madison, WI, USA) and the leftover was ethanol precipitated. The success of rRNA removal was determined by measurement on TapeStation using high-sense RNA ScreenTape and Reagents (all from Agilent, Santa Clara, CA, USA). Next, RNA was purified and fragmented; first strand cDNA synthesis was performed using SuperScript II (Thermo Fisher Scientific, Waltham, MA, USA) followed by second strand cDNA synthesis, end repair, 3’-end adenylation, adapter ligation and PCR amplification. All of the purification steps were performed using AmPureXP Beads (Beckman Coulter, Indianapolis, IN, USA). Final libraries were quality checked using D1000 ScreenTape and Reagents on TapeStation (all from Agilent, Santa Clara, CA, USA). The concentration of each library was determined using the KAPA Library Quantification Kit for Illumina (KAPA Biosystems, Wilmington, MA, USA). Sequencing was performed on an Illumina NextSeq instrument using the NextSeq 500/550 High Output Kit v2 (300 cycles; Illumina, San Diego, CA, USA) generating ~10 million clusters for each sample.

### Bioinformatic analysis of RNA-sequencing data

Following sequencing, raw paired-end Illumina reads were quality trimmed in CLC Genomics Workbench Tool (v.11.0, Qiagen bioinformatics, Aarhus, Denmark) using an error probability threshold of 0.01. No ambiguous nucleotide was allowed in trimmed reads. For filtering, reads were mapped on CRISPR spacer sequences using CLC with a length fraction of 0.9 and a sequence identity threshold of 0.95. Only those read pairs were kept for the subsequent RNA-Seq analysis that displayed no mapping against the CRISPR spacer construct. RNA-Seq analysis package from CLC was then used to map filtered reads on a custom-masked *E. coli* K12 MG1655 genome version (based on U00096.3, harbouring only one copy from each relevant IS group: IS*1*A, IS*1*F, IS*3*, IS*5*, IS*5*Y, and IS*150*). Only those reads were considered that displayed an alignment longer than 80% of the read length while showing at least 95% sequence identity against the reference genome. Next “Total gene read” RNA-Seq count data was imported from CLC into R 3.3.2 for data normalization and differential gene expression analysis. Function “calcNormFactors” from package “edgeR” v.3.12.1 was used to perform data normalization based on the “trimmed mean of M-values” (TMM) method (43). Log transformation, linear modeling, empirical Bayes moderation as well as the calculation of differentially expressed genes were carried out using “limma” v. 3.26.9, (44). Genes showing at least two-fold gene expression change with an FDR value below 0.05 were considered as significant.

### Data availability

Gene Expression Omnibus (GEO) archive of the six sequenced libraries (three IS-knockdowns and three controls) was deposited in NCBI’s GEO Archive at http://www.ncbi.nlm.nih.gov/geo under accession GSE110946.

### Plasmid availability

pCRISP_IS and pCRIS will be made available at Addgene https://www.addgene.org/. The sequence map of pCRIS is displayed in GenBank format in the Supplement.

### Measurement of chromosomal mutation rate

To infer the rate of IS transposition to chromosomal targets, we used two established methods. The first method detects mutations of the *cycA* gene causing D-cycloserine resistance and has been described earlier (16). Briefly, 20 parallel cultures of the tested strain were fully grown at 37 °C in MS+Cm and a fraction of each was plated on D-cycloserine-containing MS+Cm plates. The appropriate dilution of three random cultures was also plated on LB+Cm plates to infer the mean total cell number. The Ma-Sandri-Sarkar Maximum Likelihood method of fluctuation analysis was used to calculate the mutation per culture (m) based on the numbers of cycloserine-resistant colonies (45). The m values of strains assayed in parallel were compared using unpaired, two-tailed t-tests, as described previously (46). To obtain and display a biologically more relevant measure of mutations, m was divided by the total cell number to yield the mutation rate (mutation/gene/generation) of the *cycA* gene. The m values were not accepted to be significantly different for two strains if their total cell numbers were also significantly different, or if the mutation rates and the m values displayed an opposite pattern. PCR-amplification of the *cycA* locus using primers cycA1 and cycA2 allowed the classification of mutants as point mutants, deletion mutants or insertion mutants based on amplicon length. Further PCR-assay of the insertion mutants using IS-specific primers (**Table S1**) yielded information on the contribution of each IS type to the disruption of the *cycA* gene.

The second method to monitor IS transposition measures the spontaneous activation of the cryptic *bgl* operon in cells starving on salicin-minimal plates (23). Cells to be assayed were grown to saturation in 20 parallel 1-ml cultures at 37 °C in MS+Cm medium. Cells of each tube were pelleted, and plated onto MS+Cm plates containing 0.4 % salicin (Alpha Aesar, Ward Hill, MA, US) as the sole carbon source, and incubated at 37 °C. The appearing colonies were counted and logged daily for ten days. Total cell counts were obtained by plating appropriate dilutions onto LB+Cm plates. Statistical comparison of parallel lines was carried out in two ways: i) Mutation frequencies were calculated by dividing the mean colony numbers observed on day 10 by the total cell count. Mutation frequencies were compared using Mann-Whitney U-tests. ii) Total cell-normalized mean daily increments of mutant numbers were calculated for control and IS-silenced strains and compared using unpaired, two-tailed t-tests.

Classification of mutants as insertion-mutants or “other” types of mutants was based on the amplicon-length of the PCR-amplified *bgl* region using primers bglR1 and bglR2 (**Table S1**). Insertion mutants were called when PCR fragments were longer than amplicons obtained from wild type cells. PCR analysis of insertion mutants using IS-specific and *bgl*-specific primers allowed the identification of the IS-type integrating into the *bgl* regulatory region (*bglR*).

### Adaptive laboratory evolution to measure plasmid-stability

*E. coli* DH5αZ1 or JM107MA2 cells carrying the pBDP_Km_GFP5 plasmid were transformed either with pCRISPathBrick (serving as the control) or pCRIS. A single colony from each transformation was inoculated and was fully grown in LB+Ap+Cm medium as a starter culture, diluted 1000-fold in fresh LB+Ap+Cm+IPTG medium and divided each into 48 parallel cultures of 200 μl. These cultures were shaken in 96-well microplates (Greiner Bio-One International, Kremsmünster, Austria) in a SynergyHT microplate reader (BioTek, Winooski, VT, USA) at 37 °C, and supplemented with Km after 2 h. After overnight shaking and growth, the cultures were diluted 400-fold into fresh LB+Km+Cm+IPTG during their transfer to a new microplate. Such growth and 400-fold dilution was repeated 3–7 more times. During growth, OD_600_ and GFP-fluorescence readings (excitation filter: 485/20, emission filter: 528/20) were recorded at 15-minute intervals. The experiments were evaluated two ways: A) The mean fluorescence of both cell lines was calculated every day from the daily peak GFP fluorescence values detected in each corresponding well, and was compared for the two lines using unpaired, two-tailed t-tests. B) Every day, each well was classified as ‘active’ or ‘inactive’ depending on whether the peak green fluorescence intensity reached a certain threshold value (3000 AU) or not, respectively, and the number of ‘inactive’ wells was compared for the two strains using χ ^2^ tests.

To assess the type of mutations that were present after the adaptive evolution, we performed a PCR-based assay. PCR amplification of the VF2-VR segment (comprising the *km* and *gfp* genes) directly from the cultures allowed the identification of insertion mutants based on increased product length. Then, a second round of PCR was carried out on insertion mutants using primer VF2 and various IS-specific primers (**Table S1**) to permit the identification of the exact IS types residing in the expression cassette.

## RESULTS AND DISCUSSION

### Simultaneous silencing of up to 38 mobile genetic elements in *E. coli*

To develop a portable method that can downregulate insertional mutagenesis genome-wide we established a plasmid-encoded CRISPRi system that targets selected transposases. Specifically, we targeted an extrachromosomally expressed, catalytically inactive SpCas9 (dCas9) to the promoter sequence in the left inverted repeat of selected mobile genetic elements in *E. coli*. We chose IS*1*, IS*3*, IS*5* and IS*150* to be targeted by our construct due to their main contribution to insertional mutagenesis in *E. coli*, as demonstrated by two previous analyses (16,23).

We hypothezised that our system allows the inhibition of transposition by a) transcriptional silencing of the transposase gene, and b) limiting the access of the transposase enzymes to the left inverted repeat of the mobile element and thereby inhibiting DNA cleavage and subsequent transposition. We implemented the well-established and robust pCRISPathBrick plasmid set that i) constitutively expresses the dCas9, as well as one or more crRNAs of choice, ii) permits CRISPR array assembly with a relative ease and iii) has been applied previously for multi-target silencing *in vivo* (34).

Spacers targeting these IS elements were designed, synthesized, and cloned into the pCRISPathBrick plasmid to construct pCRIS (see Materials and Methods). In this way, a total of 26 IS elements were targeted simultaneously by CRISPRi in *E. coli* K-12 MG1655, and 38 copies in BL21(DE3), as listed in **Table 1**.

**Table 1.** Copy numbers of relevant IS elements in *E. coli* strains MG1655 and BL21(DE3)

|  | MG1655 | BL21(DE3) |
| --- | --- | --- |
| IS1 | 8 | 29 |
| IS3 | 5 | 4 |
| IS5 | 12 | 0 |
| IS150 | 1 | 5 |
| **Total** | **26** | **38** |

### Genome-wide simultaneous suppression of transposases from pCRIS

First, we investigated whether the expression of CRISPRi from a single episomal vector (pCRIS) is capable of suppressing transposases at multiple chromosomal loci. To this aim, the effect of the pCRIS plasmid on the host transcriptome was compared to that of a control plasmid using a standard Illumina RNAseq assay (see Materials and Methods) and the expression of each gene was quantified as the frequency of Illumina sequencing reads that map to the given target (**Figure S1-S3**).

Considering changes in transposase expression, the presence of pCRIS resulted in a more than four-fold silencing of IS*1*A (t-test: p<8×10^-8^), compared to a non-suppressed control (**Fig. 1A**). Moreover, the first open reading frame of IS*1*F (gene b4294) showed an even stronger, 11-fold suppression (t-test: p<2×10^-4^) (**Fig. 1B**). Dramatic knockdown was observed for the transposases of both the IS*3* and the canonical IS*5* elements, displaying more than 360- and 26-fold reduction of sequencing-read frequencies, respectively (p<2×10^-5^, p<2×10^-14^, respectively)(**Fig. 1C, D**). Interestingly, no transcriptional knockdown was observed for IS*150* (Supplementary **Fig. S4**). The lack of changes in IS*150* mRNA levels may indicate the necessity to re-define the active promoter of this element. This issue, along with other features of the transcriptomic dataset are discussed in Supplementary Note 1.

**Figure 1.**
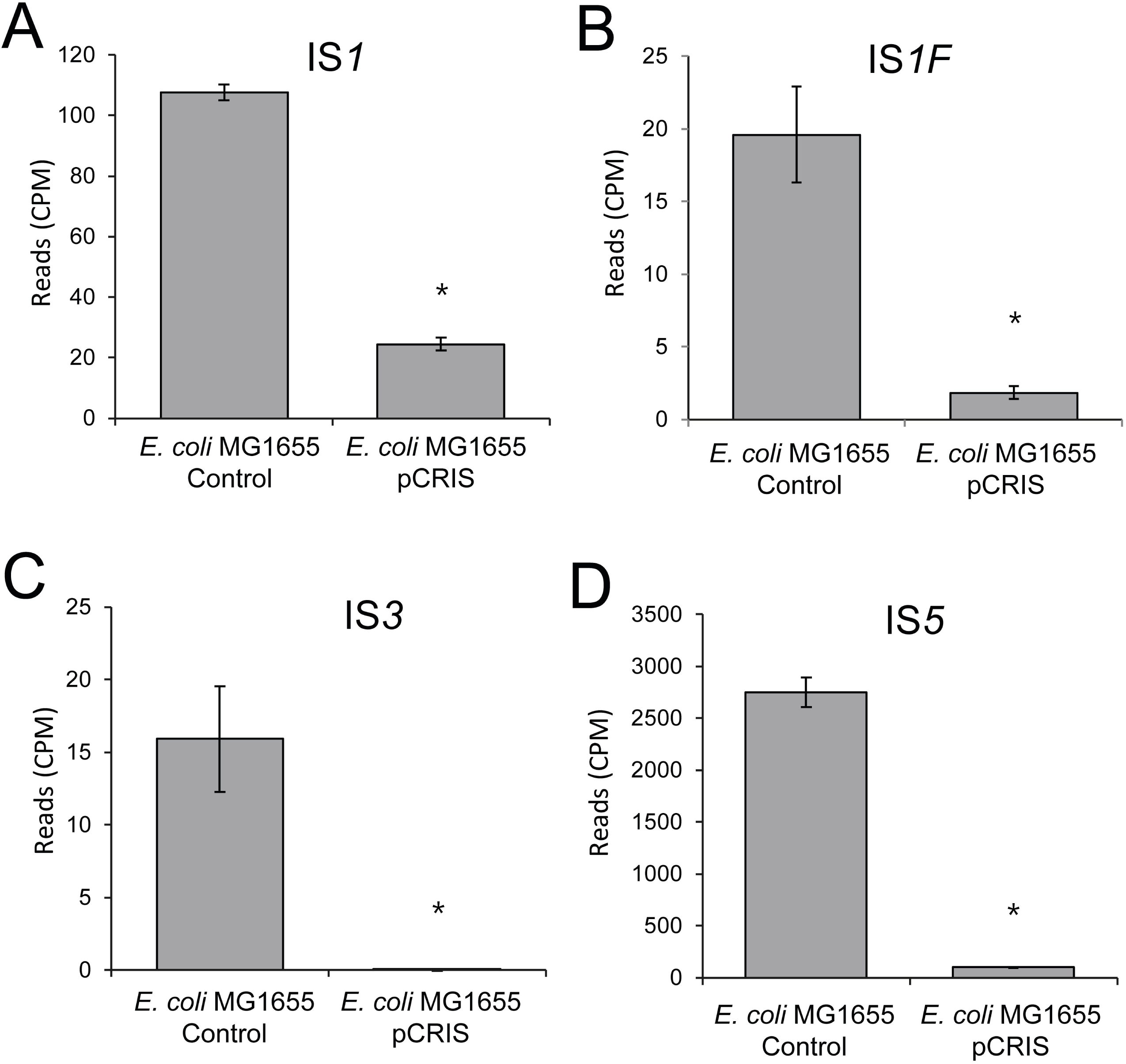
Transcriptional changes of IS elements in *E. coli* K-12 MG1655 caused by the propagation of the pCRIS plasmid. The figure displays the frequency of Illumina sequencing reads mapping to IS*1* (A), IS*1F* (B), IS*3* (C) and IS*5* (D). Star denotes significance (p<2×10^-4^) based on t-test. Error bars represent SD, *n* = 3. CPM: counts per million reads

### pCRIS efficiently suppresses chromosomal insertional mutagenesis

Next, we assessed the effect of CRISPRi on the spontaneous chromosomal mutational landscape of *E. coli*. We sought to precisely evaluate insertional mutagenesis caused by endogenous insertional sequences, primarily IS*1*, IS*5*, IS*3*, and IS*150*. For this aim, we applied two complementary mutation-detection systems. The first experimental setup detects deleterious mutations of the *cycA* gene. CycA, which is responsible for D-cycloserine susceptibility, can be inactivated by various loss-of-function mutations, including IS transposition (16,23). Based on the numbers of D-cycloserine-resistant colonies observed, we carried out fluctuation analyses to compare the rate of mutations inactivating *cycA* in the presence and the absence of pCRIS-based transposase-suppression (see Materials and Methods). In our tests, the mutation/culture values (*m*) were significantly lower in *E. coli* K-12 MG1655 in the presence of pCRIS compared to a non-targeting plasmid, pCRISPathBrick (**Fig. 2A**). Notably, the observed effect of pCRIS-based IS-silencing on the mutation rate was comparable to that caused by the complete genomic deletion of ISes (23) which suggests the near-complete elimination of insertional mutagenesis.

**Figure 2.**
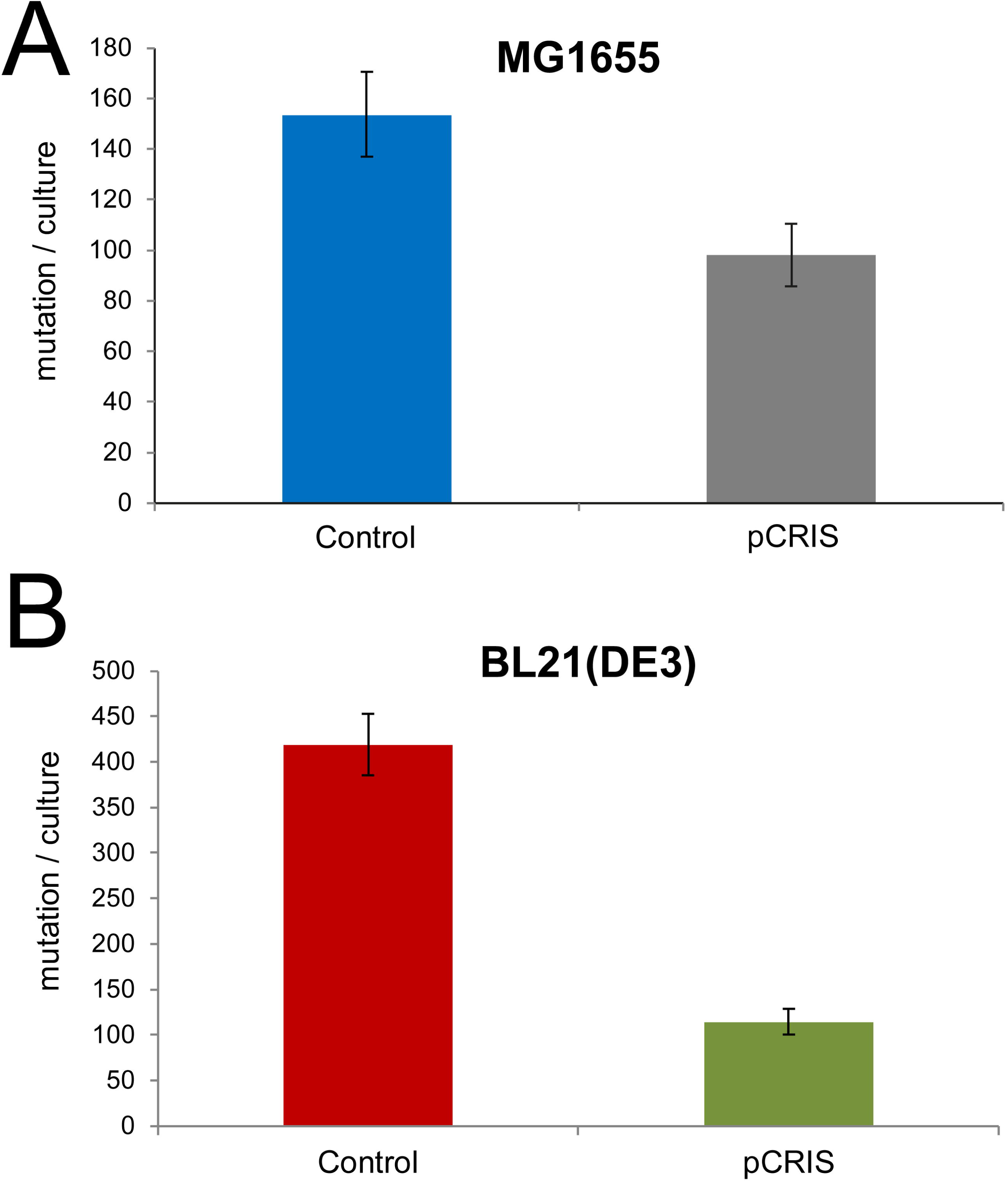
Effect of pCRIS-based IS-knockdown on the rate of *cycA* mutations in multiple *E. coli* strains: (A) *E. coli* K-12 MG1655, (B) *E. coli* BL21(DE3). Error bars represent SD based on n=3 independent measurements. * indicates significance (p<0.05) based on t-test.

Strikingly, the reduction of insertional mutagenesis caused by pCRIS was even more remarkable in the industrial *E. coli* host, BL21(DE3). Applying the same assay, a 73% decrease in *m* was observed as compared to the control (**Fig. 2B**). We attributed this large effect to the high fraction of IS*150* insertions among *cycA* mutants in BL21(DE3), seen earlier (24). Confirming our hypothesis, PCR analysis of 100 *E. coli* BL21(DE3) *cycA* mutants indicated that in the non-suppressed, parental strain, 78% and 7% of the mutations are caused by the transposition of IS*150* and IS*1*, respectively (**Fig. 3A**). However, in pCRIS-harbouring *E. coli* BL21(DE3) cells only 1% of the resistant variants displayed IS integration with the rest being point mutants. Moreover, no IS*1* transposition could be detected among 100 tested colonies (**Fig. 3B**). Therefore, the large decrease in mutation events seen in *E. coli* BL21(DE3) in the presence of pCRIS (**Fig. 2B**) can be attributed to the near-complete elimination of IS transposition into *cycA* (**Fig. 3**). The changes observed in the composition of mutants caused by the presence of pCRIS also underline the fact that *IS*150 mobility was dramatically downregulated despite the lack of transcriptional silencing.

**Figure 3.**
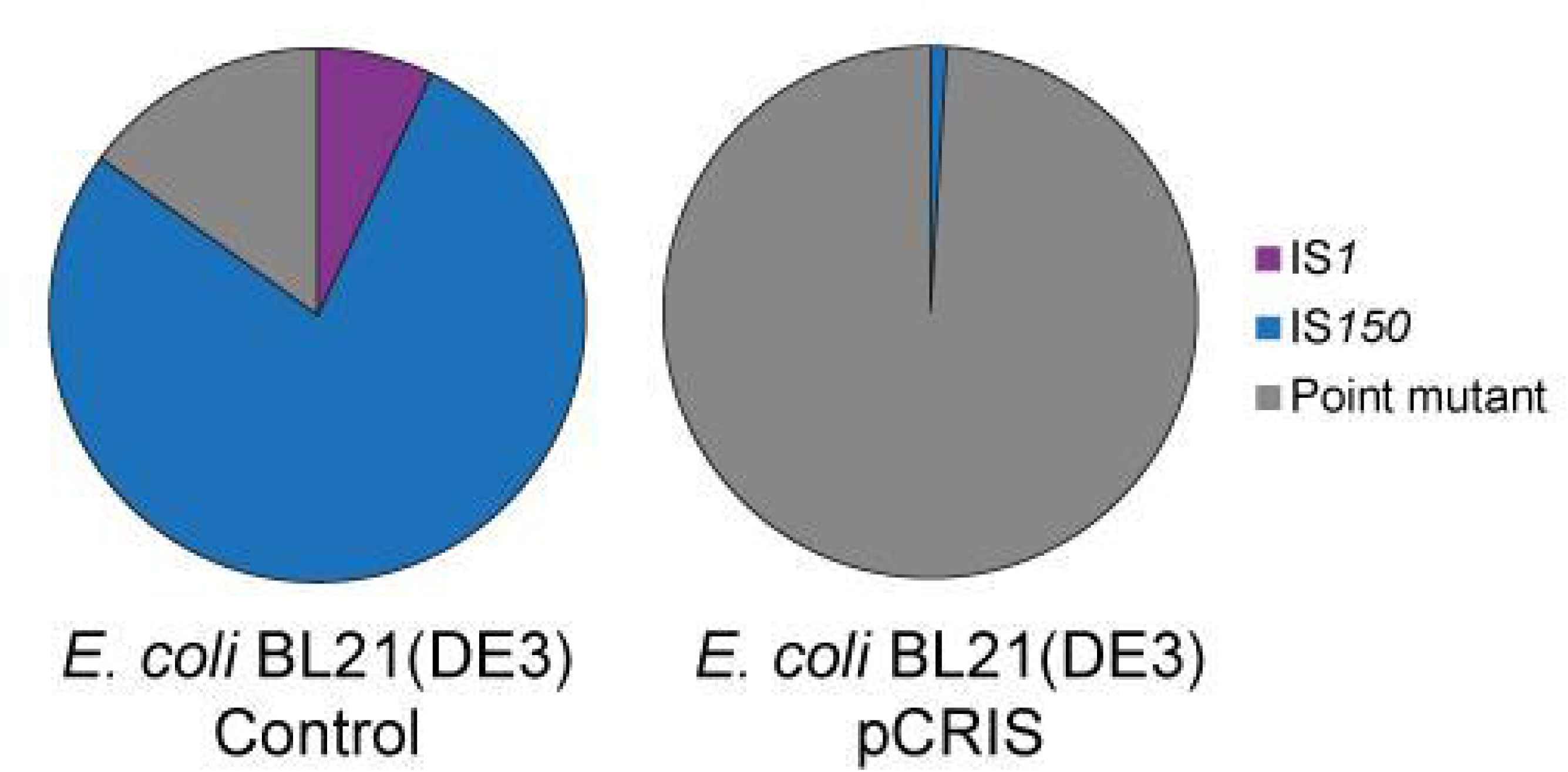
Effect of IS-knockdown on the spectrum of mutations inactivating *cycA* in *E. coli* BL21(DE3). (A) Effect of a non-targeting control plasmid (pCRISPathBrick); (B) effect of pCRIS-based transposase-silencing.

The second mutagenesis assay that we applied tests the activation of the cryptic *bgl* operon in *E. coli* that leads to the ability of mutants to grow on salicin-minimal medium. This assay was chosen for three noteworthy features: First, it investigates insertional mutagenesis at a locus distinct from *cycA*. Second, it is highly specific for IS transposition (17): in case of wild-type *E. coli* K-12 MG1655, more than 90% of salicin-assimilating colonies contain an IS element (primarily *IS1* and IS*5*) at the *bglR* locus. And third, besides detecting mutations in growing cultures, it is also capable of quantifying mutation events in non-growing living cells, starving on salicin minimal medium.

When comparing pCRIS-containing *E. coli* K-12 MG1655 cells to controls carrying a non-targeting plasmid, this assay revealed a 4.4-fold decrease in overall mutation frequency in stationary phase (**Fig. 4**). PCR-analysis of the salicin-utilizing colonies after 10 days uncovered that the frequency of insertion mutants decreased nearly 11-fold, with the near complete elimination of IS1-transpostion into *bglR* (**Fig. 4**). The daily rate of the emergence of salicin-assimilating cells during the 10-day stationary phase also displayed a five-fold reduction in the presence of pCRIS (p<10^-3^) (**Fig. S5**).

**Figure 4.**
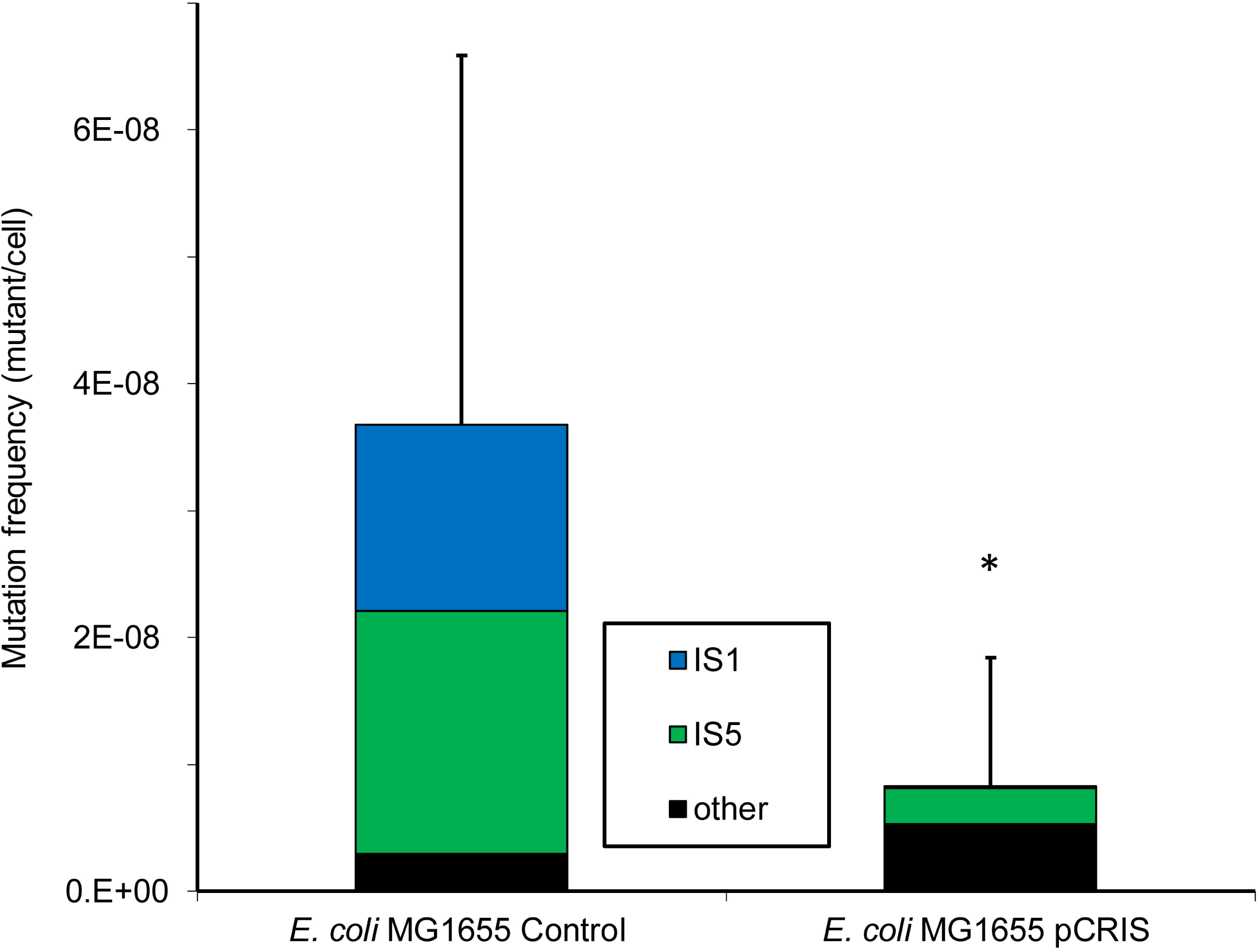
Effect of IS-knockdown on the frequency and spectrum of mutations activating the *bgl* operon of *E. coli* MG1655. The frequency and composition of salicin-assimilating mutants observed on day 10 are shown. The control strain carried the non-targeting pCRISPathBrick plasmid, pCRIS represents active IS-silencing. * p<10^-4^ (Mann-Whitney U-test).

Based on these two assays detecting mutation events at two distinct chromosomal loci, IS-knockdown from a single plasmid was found to efficiently downregulate insertional mutagenesis in two different strains of *E. coli*.

### Transposase suppression increases the genetic stability of plasmid-borne synthetic DNA

Finally, we sought to investigate the effects of transposase-suppression on the genetic stability of plasmid-borne synthetic DNA elements during their long-term propagation in *E. coli*. To estimate the genetic stability of a synthetic DNA construct under mock-industrial conditions, we carried out adaptive evolution experiments utilizing a previously established plasmid-based system that is reportedly susceptible to insertional mutagenesis (7).

Specifically, our test plasmid pBDP_Km_GFP5 carries a *gfp* gene and an oppositely oriented kanamycin-resistance gene, driven by a bi-directional LacI-repressible promoter (7), called the *km-gfp* cassette. In its induced state kanamycin-selection transcriptionally links GFP expression to the host cell’s survival and thereby prevents mutations that disrupt the bi-directional promoter. As a result, the burden of GFP production is most often relieved by the IS-mediated interruption of its gene, instead of the otherwise commonly seen promoter mutations. In our evolutionary experiments, continuously-induced pBDP_Km_GFP5 was propagated in the LacI-overexpressing DH5αZ1 host in 48 parallel cultures for 60–65 generations in the presence of kanamycin. Notably, the timeframe and the number of cell divisions in these experiments were industrially relevant as cell-growth from a seeding starter to a fermentation volume of 1 – 200 m^3^ requires approximately 60 generations. The effect of IS-knockdown was assessed by comparing pCRIS-carrying strains with control lines harbouring the non-targeting pCRISPathBrick plasmid. To evaluate mutational dynamics in the course of the adaptation we continuously monitored GFP expression and the OD_600_ value of each culture and thereby assessed the integrity of the synthetic contsruct (Materials and Methods).

The means of the maximal intensities were calculated each day for each strain to follow the day-to-day changes in the average heterologous protein production (**Figure S6**). By the end of laboratory evolution, the median fluorescence of the bacterial populations dropped by 16% when pCRIS was present in cells, in contrast to the 35% decrease without controlling insertional mutagenesis (p<2×10^-3^)(**Fig. 5)**. Similar results were obtained when repeating the experiment in *E. coli* JM107MA2 (**Figs. S7, S8**).

**Figure 5.**
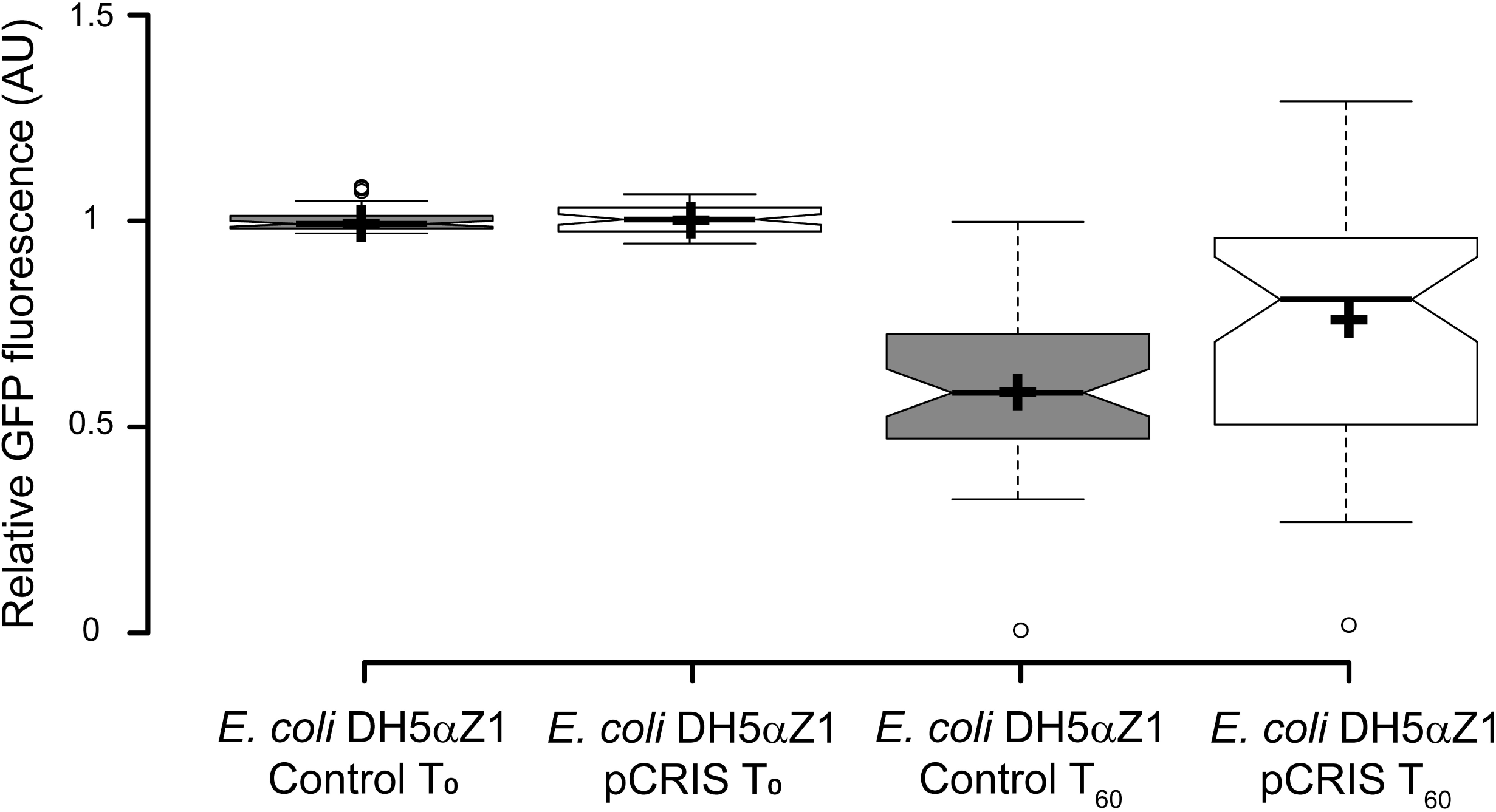
The effect of IS-silencing on plasmid stability in *E. coli* DH5αZ1. The figure shows the T0-normalized, relative GFP-signal (AU) of 48 parallelly passaged cultures of *E. coli* DH5αZ1 that are either carrying the non-targeting, control plasmid (pCRISPathBrick) or the IS-knockdown plasmid, pCRIS. Fluorescence intensities were compared on initiation (T0) and after 60 generations (T60) in the course of adaptive laboratory evolution. Center lines show the medians; box limits indicate the 25th and 75th percentiles as determined by R software; whiskers extend 1.5 times the interquartile range from the 25th and 75th percentiles, outliers are represented by dots; crosses represent sample means. n=48 sample points. * marks the significant difference of the means (p<2×10^-3^) assessed by a Mann-Whitney U-test.

To investigate the mutational processes and dynamics behind synthetic construct inactivation we monitored the transfer-to-transfer changes observable in each individual culture in the course of our laboratory adaptive evolution experiment. Each well was classified as ‘inactive’ or ‘active’ based on its maximal GFP fluorescence intensity and the number of ‘inactive’ cultures was compared for the two strains. After 7 days and 60 generations of serial passage (T60), only one ‘inactive’ culture was found for pCRIS while the non-suppressed control inactivated eleven cultures, thus indicating a significant decrease in the frequency of inactivating mutations (χ ^2^ test: p<0.005) (**Figure 6A**).

**Figure 6.**
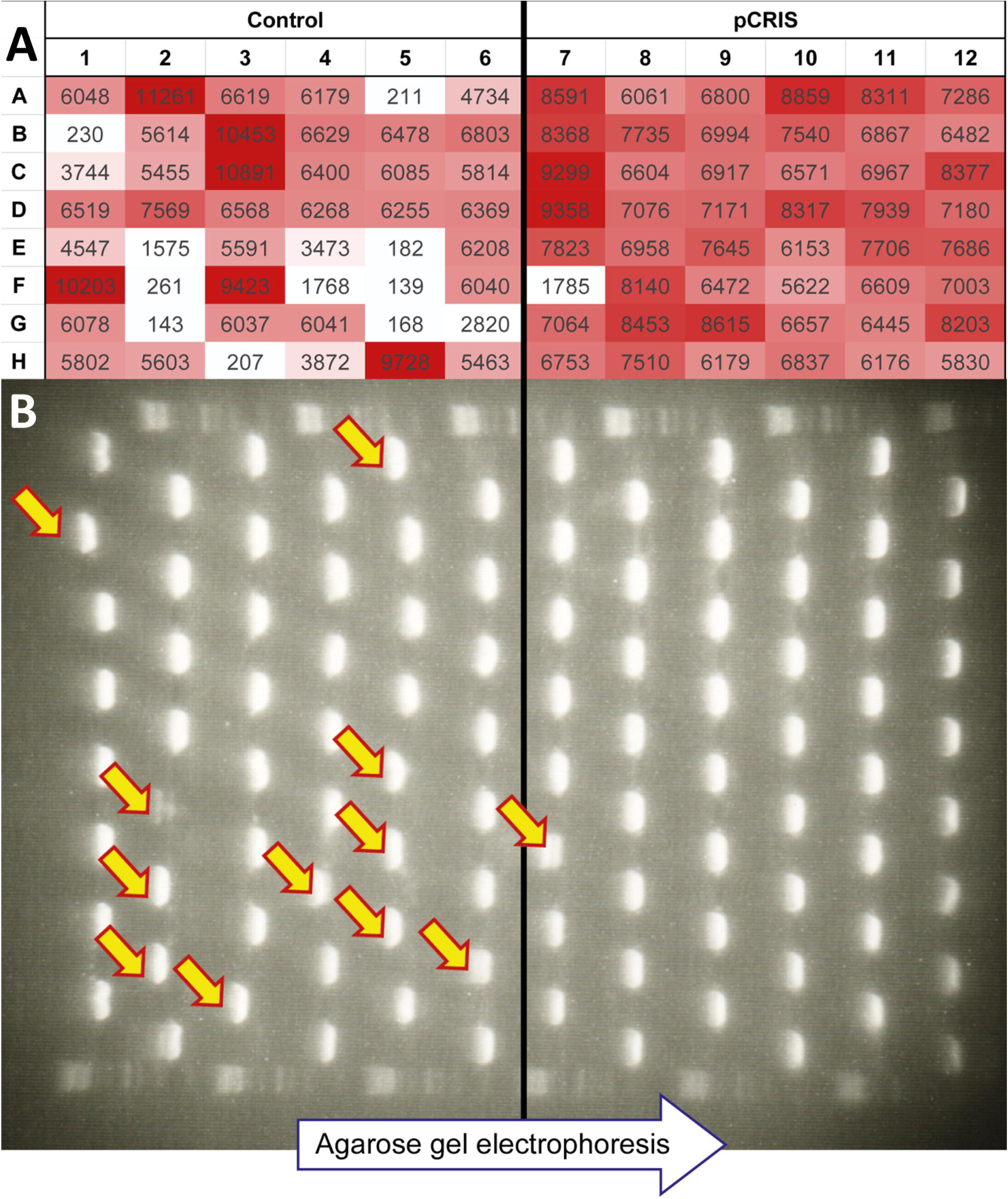
Effect of IS-knockdown on fluorescence of individual cultures. (A) Maximum fluorescence intensities (A.U.) of the individual cultures in the presence of either an IS-silencing plasmid (pCRIS) or a non-targeting control plasmid (pCRISPathBrick), after 7 days (60 generations) of continuous propagation (T60). (B) Analysis of the mutagenic spectrum within the *km-gfp* cassette of pBDP_Km_GFP5 from individual cell-populations (n=48) at T60. Arrows mark PCR products displaying an increased length due to IS-transposition into *km-gfp*.

Next, we sought to investigate the causative mutations behind inactivation. PCR-amplification of the *km-gfp* cassette demonstrated that the dominant genotype in each ‘inactive’ culture corresponded to an elongated PCR product, thereby indicating that each inactivating mutation was caused by IS transposition into pBDP_Km_GFP5 **(Fig. 6B)**. Further PCR-based analysis of these insertion mutants in the non-suppressed control strain revealed that 10 out of 11 was caused by IS*2* transposition, with IS*10* being responsible for the remaining one. The single insertion seen in the IS-silenced strain was caused by IS*10*. The repetition of our adaptive laboratory evolution in a further strain of *E. coli* (JM107MA2) revealed a smaller, but significant stabilizing effect for pCRIS (**Figs. S7, S8**). The significant suppression of IS*2* transposition, albeit not its complete elimination was observed in JM107MA2, as well (**Table S4**.). Since our CRISPR array did not target IS*2*, this observation was surprising. The possible background of this intriguing phenomenon is discussed in Supplementary Note 2.

In sum, these results demonstrated that continuous IS-knockdown is capable of extending the half-life of episomally propagated synthetic genetic constructs under conditions that are similar to industrial fermentations. We expect that pCRIS can offer a portable alternative to time-consuming, genome-wide elimination of insertional sequences (23,24) where increased genetic stability is required. However, follow-up studies will be needed to evaluate the effects of pCRIS on industrially-applied biosynthetic pathways and under fermentation conditions.

## DISCUSSION

One of the greatest challenges limiting synthetic biology is the fact that biological systems constantly evolve, and the direction of evolution seldom matches that desired by bioengineers (1). Decelerating evolution by limiting mutational processes provides a promising strategy to increase genetic stability, however the genome-scale elimination of transposition has, to date lacked a simple and generally applicable solution. Our work addresses this problem by introducing a portable system, termed pCRIS that reduces insertional mutagenesis within multiple *E. coli* strains and circumvents the time-consuming nature of previous genome-engineering endeavors. pCRIS displays several unique features, it is: a) portable, demonstrated by its application in distinct *E. coli* strains; b) instantaneous, requiring only a simple transformation into the strain-of-interest; c) reversible, a plasmid curing is sufficient to regain wild-type mutation rate in the host; and d) highly specific for insertional mutagenesis.

Several examples are available in the literature to limit the rate of single-nucleotide exchanges or homologous recombination-driven rearrangements in bacterial hosts by inactivating a handful of genes (8–13). Such a simple and rapid solution to counteract transposable elements has been lacking, despite increasing evidence concerning their role in transgene-inactivation. For example, Rugjberg et al. (14), investigated the spectrum of mutations inactivating industrially-applicable mevalonate-producing pathways. Their results showed a striking dominance of insertions with IS transposition being responsible for 69.5 to 99.9 % of mutations in *E. coli* DH10B. Moreover, a significant number of case studies from basic- and applied-research have also reported the interruption of cloned DNA segments by IS elements (47–53).

In this work, we demonstrate the CRISPRi-based knockdown of IS-activities in multiple *E. coli* strains, including the biotechnologically useful production host, *E. coli* BL21(DE3). The system, termed pCRIS, efficiently suppressed the mobility of four types of IS elements (IS*1*, IS*3*, IS*5*, and IS*150*), that notably distinguishing it from other endeavours (54). By expressing a single array of targeting constructs that directs a catalytically inactive SpCas9 to the transposases-of-interest, IS-transposition to chromosomal and episomal targets was demonstrated to be significantly downregulated in various experimental setups. Importantly, the expression of less than 0.3% of all *E. coli* genes differed upon IS-knockdown, which demonstrates the unique specificity of plasmid-based IS suppression (Supplementary Figure S3). Albeit multiplex gene-silencing has been established before using the same approach (34), to the best of our knowledge, this is the first time multiple crRNAs each targeting multiple loci have been used effectively to control transcription. We demonstrate this ‘multi-multi’ strategy to successfully control the mobility of up to 38 copies of IS elements in *E. coli* BL21(DE3). Most notably, pCRIS, while requiring only a single plasmid delivery performed within a single day, provided a reduction of IS-mobility comparable to that seen in genome-scale chromosome engineering projects (23,24). It may thereby provide rapid strain stabilization for a wide range of applications in basic- and applied-research while circumventing the time-consuming nature of genome-scale engineering.

However, our system also has some limitations, as the current design of pCRIS may not necessarily be the optimal choice for all projects that require IS-suppression. First, the prevalence of IS-types found in inactivated mutants shows dependence on the target sequence, the host strain, and possibly other factors as well, which may require modification of the plasmid. The user-friendly cloning system of pCRISPathBrick, however allows rapid cloning of repeat-spacer tandems targeting further mobile elements (34). Second, pCRIS is based on a p15A origin of replication that can be incompatible with several other production-plasmids and may be restricted to certain hosts. The use of broad-host replication origins, combined with modular plasmid assembly strategies can provide solutions for such issues for a wide the range of applications (55,56). Third, the constitutive overexpression of dCas shown here can itself cause a negative fitness effect on the host, which may prove to be deleterious when combined with the overexpression of a toxic gene of interest. Such cases may necessitate the fine-tuning of dCas9 using inducible promoters, or require the chromosomal integration of the CRISPR/dCas machinery. And finally, certain plasmid-based expression systems are not interrupted by insertions at all, but display other modes of inactivation, such as deletions caused by replication slippage (7). Further solutions, discussed in the introduction must therefore always be kept in mind.

In summary, pCRIS increases genetic stability of chromosomal- and plasmid-based genetic circuits via the introduction of a single plasmid. The specific nature of IS-silencing may make pCRIS useful in a limited number of hosts, we nevertheless expect that the robustness of CRISPRi will permit our system to be easily reprogrammed to target other IS types and thereby yield a full set of strain-specific IS-silencing plasmids. With such possibilities in mind, we expect that the rationale of pCRIS will be broadly applicable to decelerate transposition-driven evolutionary processes in a wide variety of laboratory model-strains and industrially applied bacterial hosts.

## FUNDING

This work was supported by the National Research, Development, and Innovation Office of Hungary (NKFIH) Grant No. K119298 (FT) and the GINOP-2.3.2-15-2016-00001 (FT) and by grants from the European Research Council H2020-ERC-2014-CoG 648364 - Resistance Evolution (to C.P.), the Wellcome Trust (to C.P.), and GINOP (MolMedEx TUMORDNS) GINOP-2.3.2-15-2016-00020, GINOP (EVOMER) GINOP-2.3.2-15-2016-00014 (to C.P.),; the ‘Lendület’ Program of the Hungarian Academy of Sciences (to C.P.); and a PhD fellowship from the Boehringer Ingelheim Fonds (to Á.N.).

## ACKNOWLEDGMENTS

We thank Prof. Luciano Marraffini and Prof. Herbert M. Sauro for providing plasmids.*Conflict of interest statement*. None declared.

